## Supplementary Table 1 for "Fetal exposure to endocrine disrupting-bisphenol A (BPA) alters testicular fatty acid metabolism in the adult offspring: relevance to sperm maturation and quality"

**Supplementary Table 1**: Predesigned SYBR green I rat primers and corresponding genes used for the mRNA expression analyses

| **Sl.**  **no.** | **Primer ID** | **Gene symbol** | **Gene ID** | **Gene name** | **Nucleotide sequences (5’-3’)** | **Ref_seqID** |
| --- | --- | --- | --- | --- | --- | --- |
| 1 | R1_Fabp9 | FABP9 | [64822](https://www.ncbi.nlm.nih.gov/gene/64822) | Fatty acid-binding protein 9 | F 5'- TAGCATTAGTTTCAATGGGG-3'  R 5'- GTTATAAGGCTCTTCACTTTCC-3' | [NM_022854](http://www.ncbi.nlm.nih.gov/entrez/query.fcgi?db=Nucleotide&cmd=Search&term=NM_022854&doptcmdl=GenBank) |
| 2 | R1_Cox2 | COX2 | 29527 | Cyclooxygenase 2 | F 5'- CTCATACTGATAGGAGAGACG-3'  R 5'- TCGAACTTGAGTTTGAAGTG-3' | NM_017232 |
| 3 | R1_Acsbg2 | ACSBG2 | [301120](https://www.ncbi.nlm.nih.gov/gene/301120) | Long chain fatty acid-CoA ligase | F 5'- CATGACAACATCACATGGAC-3'  R 5'- TTTGATAGGGATCCAGATGTC-3' | [NM_001080096](http://www.ncbi.nlm.nih.gov/entrez/query.fcgi?db=Nucleotide&cmd=Search&term=NM_001080096&doptcmdl=GenBank) |
| 4 | R1_Lpl | LPL | 24539 | Lipoprotein lipase | F 5'- CCTACTCCTTCTTGATTTACAC-3'  R 5'- GAAGATGACCTTTTTCTGAGTC-3' | NM_012598.2 |
| 5 | R1_Lipe | LIPE | 25330 | Lipase E, hormone sensitive type | F 5'- GTGGAAAGATGTCAGGATATG-3'  R 5'- GTAAATCCATGCTGTGTGAG-3' | NM_012859 |
| 6 | R1_Slc25a20 | SLC25A20 | 117035 | Carnitine-acylcarnitine translocase | F 5'- AGCCCACCTGTTATCCACTG-3'  R 5'- TGTGCAAAAAGAGCCTTCCT-3' | NM_053965.2 |
| 7 | R1_Catsper2 | Catsper2 | [366174](https://www.ncbi.nlm.nih.gov/gene/366174) | Cation channel sperm-associated protein 2 | F 5'- TGTTGCTTGGTTCCATTATC-3'  R 5'- TTGACTGGTTCCTCTTAGTG -3' | [NM_001012220](http://www.ncbi.nlm.nih.gov/entrez/query.fcgi?db=Nucleotide&cmd=Search&term=NM_001012220&doptcmdl=GenBank) |
| 8 | R1_Catsper1 | Catsper1 | 689349 | Cation channel sperm-associated protein 1 | F 5'- AAACCCATCACCACTATGAG-3'  R 5'- CCAGATCTTTCCTGGTTTTG-3' | XM_001070492 |
| 9 | R1_Fads1 | FADS1 | 84575 | Fatty acid destaurase 1 | F 5'- GTACTTCTTCTTGATTGGAC-3'  R 5'- GTAAGTGAAGAAGACACGAAC-3' | NM_053445.2 |
| 10 | R1_Fads2 | FADS2 | 83512 | Fatty acid desaturase 2 | F 5'- CTTCTTCAATGACTGGTTCAG-3'  R 5'- CTTCAGTGAACTCACAATGTC-3' | NM_031344.2 |
| 11 | R1_Elovl2 | ELOVL2 | 498728 | Fatty acid elongase 2 | F 5'- CTTGTGGTCAAAGCTTCTTC-3'  R 5'- GAGGTATTTCTTCCACCAAAG-3' | NM_001109118.1 |
| 12 | R1_Elovl5 | ELOVL5 | 171400 | Fatty acid elongase 5 | F 5'- TTCTTCGTAAGAACAACCAC-3'  R 5'- ATAGTACGAGTACATGAGGAC-3' | NM_134382.2 |
| 13 | R1_Degs1 | DEGS1 | 58970 | Delta 4 desaturase, sphingolipid 1 | F 5'- ATCTTAGCGAAGTATCCAGAG-3'  R 5'- CAGAGTCATGGAATGGTTAAG-3' | NM_053323.2 |
| 14 | R1_Scd-2 | SCD 2 | 83792 | Stearoyl-Coenzyme A desaturase 2 | F 5'- TCCAGAGGAGGTATTACAAG-3'  R 5'- CTGTTTACAAACGTCTCACC-3' | NM_031841.2 |
| 15 | R1_Scd-1 | SCD 1 | 246074 | Stearoyl-Coenzyme A desturase 1 | F 5'- ATGAGAGAAGATATCCACGAC-3'  R 5'- AGTAAAATATCCCCCAGAGC-3' | NM_139192.2 |
| 16 | R1_Fasn | FASN | 50671 | Fatty acid synthase | F 5'- AAAAGGAAAGTAGAGTGTGC-3'  R 5'- GACACATTCTGTTCACTACAG-3' | NM_017332.2 |
| 17 | R1_Igf1 | IGF1 | 24482 | Insulin like growth factor 1 | F 5'- GCACCTCCAATAAAGATACAC-3'  R 5'- TGGGCTTGTTGAAGTAAAAG-3' | NM_001082479 |
| 18 | R1_Lep | LEP | 25608 | Leptin | F 5'- CTCATCAAGACCATTGTCAC-3'  R 5'- TGAGGATCTGTTGATAGACTG-3' | NM_013076.3 |
| 19 | R1_Adipoq | ADIPOQ | 246253 | Adiponectin | F 5'- TGGCGATTTTCTCTTCATTC-3'  R 5'- AGGATTAAGAGGAACAGGAG-3' | NM_144744.3 |
| 20 | R1_Act-B | ACT β | 81822 | Actin-beta | F 5'- AAGACCTCTATGCCAACAC-3'  R 5'- TGATCTTCATGGTGCTAGG-3' | NM_031144.3 |
