## Supplementary Table 2 for "Fetal exposure to endocrine disrupting-bisphenol A (BPA) alters testicular fatty acid metabolism in the adult offspring: relevance to sperm maturation and quality"

| **Target**  **protein** | **Protein**  **name** | **Primary antibody** | **Clonality** | **Host** | **Catalog & supplier** | **Dilution** |
| --- | --- | --- | --- | --- | --- | --- |
| FADS1 | Fatty acid desaturase 1 | Anti-FADS1 | Monoclonal | Rabbit | #A0178 Abclonal | 1:2000 |
| ELOVL2 | Fatty acid elongase 2 | Anti-ELOVL2 | polyclonal | Rabbit | #A17712 Abclonal | 1:500 |
| FABP3 | Fatty acid binding protein 3 | Anti-FABP3 | Polyclonal | Rabbit | #AG25A0040 Adipogen | 1:4000 |
| FABP4 | Fatty acid binding protein 4 | Anti-FABP4 | Polyclonal | Rabbit | #PA5-30591 Thermo | 1:2500 |
| FABP5 | Fatty acid binding protein 5 | Anti-FABP5 | polyclonal | Rabbit | #A6373 Abclonal | 1:1000 |
| FABP7 | Fatty acid binding protein 7 | Anti-FABP7 | Polyclonal | Rabbit | #A11604 Abclonal | 1:1000 |
| ADRP | Adipocyte differentiation related protein | Anti-ADRP | Polyclonal | Rabbit | #PA1-16971 Thermo | 1:3000 |
| PPAR α | Peroxisome proliferator activated receptor alpha | Anti-PPAR α | Monoclonal | Mouse | #MA1-822 Thermo | 1:1000 |
| PPAR γ | Peroxisome proliferator activated receptor gamma | Anti-PPAR γ | Monoclonal | Mouse | #SC-7273 Santacruz | 1:500 |
| Actin | Beta actin | Anti-β Actin | Monoclonal | Mouse | #A5316 Sigma | 1:10000 |

**Supplementary Table 2**: The primary antibodies and their dilution used in this study
